# Chemosensation drives divergent social behavior in *Drosophila*

**DOI:** 10.1101/2024.11.05.622140

**Authors:** Binbin Wu, Erin S. Keebaugh, William W. Ja

## Abstract

The social environment can have a considerable impact on animal behavior. However, the molecular signals that drive socially sensitive behaviors are not completely defined. Using the model organism, *Drosophila melanogaster*, we investigated behavioral responses to housing status. Group housing of male flies induces robust social interactions—composed of chasing and touching—in the Dahomey strain, but not in other commonly used parental lines. Testing mixed-strain groups revealed that strain-specific variation in chemosensation, rather than in the generation of chemosignals, likely underlies the divergent group behaviors. Using locomotor activity as a higher throughput readout for increased social interactions, we found that manipulations that decrease or ablate olfaction—through surgical removal of olfactory organs, silencing of olfactory neurons, or mutations in *odorant receptor co-receptor* (*Orco*)—decrease Dahomey social sensitivity. Females of all tested strains, including Dahomey, show no evidence of increased group activity. These results demonstrate that Dahomey social sensitivity is a sexually dimorphic, olfaction-dependent group behavior. The OR43a receptor, as well as signaling through *Or43a*^*+*^ neurons, contributes to Dahomey group hyperactivity. Additional studies also suggest that strain-specific single nucleotide polymorphisms (SNPs) in the *Or43a* gene are associated with Dahomey social sensitivity. Our studies reveal insights on the mechanisms that regulate sensory recognition and behavioral responses to the social environment, and how divergent strategies may have evolved within the same species.

## Introduction

The social environment profoundly influences physiology and behavior. Negative social interactions are associated with poor health and shortened life^1,2^, highlighting the importance of understanding the molecular mechanisms that regulate social behaviors. Fruit flies (*Drosophila melanogaster*) display diverse social interactions and, coupled with the available genetic tools, provide a powerful model system for identifying the social cues and signaling pathways that drive behavior^3^.

In addition to courtship, flies exhibit collective behaviors such as movement, aggregation, and aggression that are modulated by group size, density, and sex^4,5^. Chronic isolation reduces sleep and triggers hyperphagia^6^, while chronic social grouping suppresses male-male aggression^7^. Furthermore, genetic models that capture some of the social behavioral phenotypes that accompany autism spectrum disorder (ASD) or Alzheimer’s disease (AD) have been established in *Drosophila*^8,9^. Flies deficient in Neuroligin 2, a protein associated with ASD, exhibit reduced social interactions by maintaining a greater distance from conspecifics^10^. Other studies have investigated how social environment affects fly behavior. Understanding the mechanisms that regulate group behaviors in model systems might provide insights on how social interactions are affected in various disorders.

Previous studies have implicated the role of various sensory modalities in fly social interactions, including vision, touch, and in many cases, olfaction^11–13^. In insects, cuticular hydrocarbons (CHs), synthesized by specialized cells known as oenocytes^14^, are an important class of olfactory chemosignals that mediate social interactions. CHs have complex effects on social communication due to compound characteristics such as volatility and abundance^15,16^. In *Drosophila, cis*-vaccenyl acetate (cVA) is synthesized only in males, but regulates diverse behaviors in both sexes^17^, including social group interactions^11^. Identifying chemosignals, and chemosensory mechanisms such as the specific olfactory receptor neurons (ORNs) that detect these cues to drive behavior, is crucial for understanding the strategies that animals use to modulate social behavior^18,19^.

Here, we uncovered a strain- and sex-dependent group behavior. Dahomey males show group hyperactivity that is acutely triggered by social environment and inhibited by light. We further found that group hyperactivity is dependent on olfaction, at least partially through *Or43a*^*+*^ neurons and potentially through differences in the *Or43a* gene. In humans, some social disorders are associated with olfactory defects. The presence of olfactory dysfunctions has been described as an early symptom of AD in early stages^20^, and Fragile X syndrome (FXS) patients also exhibit abnormal olfactory processing^21^. Decoding the genetic and neuronal mechanisms underlying *Drosophila* group behaviors could therefore enhance our understanding of social deficits caused by abnormal perception and processing of environmental signals.

## Results

### Dahomey males show robust social behaviors and group hyperactivity

Canton-S, *w*^1118^, and Dahomey are lines commonly used as background controls or parental strains in *Drosophila* studies. As a proxy for quantifying social interactions, solo- and group-housed flies were monitored in the Locomotor Activity Monitoring (LAM) system, which counts how often flies breach a planar array of infrared beams that bisect the chamber (Fig. 1A). At most time points, group housing suppressed the per fly activity of *w*^1118^ and Canton-S males (Fig. 1B, C). This inhibition of activity is consistent with previous studies showing that flies in groups decrease their movement to prevent aggressive contacts^22^. In contrast, group-housed Dahomey males showed robust hyperactivity at night (Fig. 1D). Group hyperactivity was acutely inhibited by light, and flies in constant darkness still showed increased but rhythmic activity per fly (Fig. S1). Introducing the *white* (*w*) mutation did not affect the parental phenotype (i.e. *w*Dahomey shows group hyperactivity; *w*Canton-S does not), enabling the use of other transgenic lines in white-eyed Dahomey or Canton-S backgrounds (Fig. S2).

**Figure 1:**
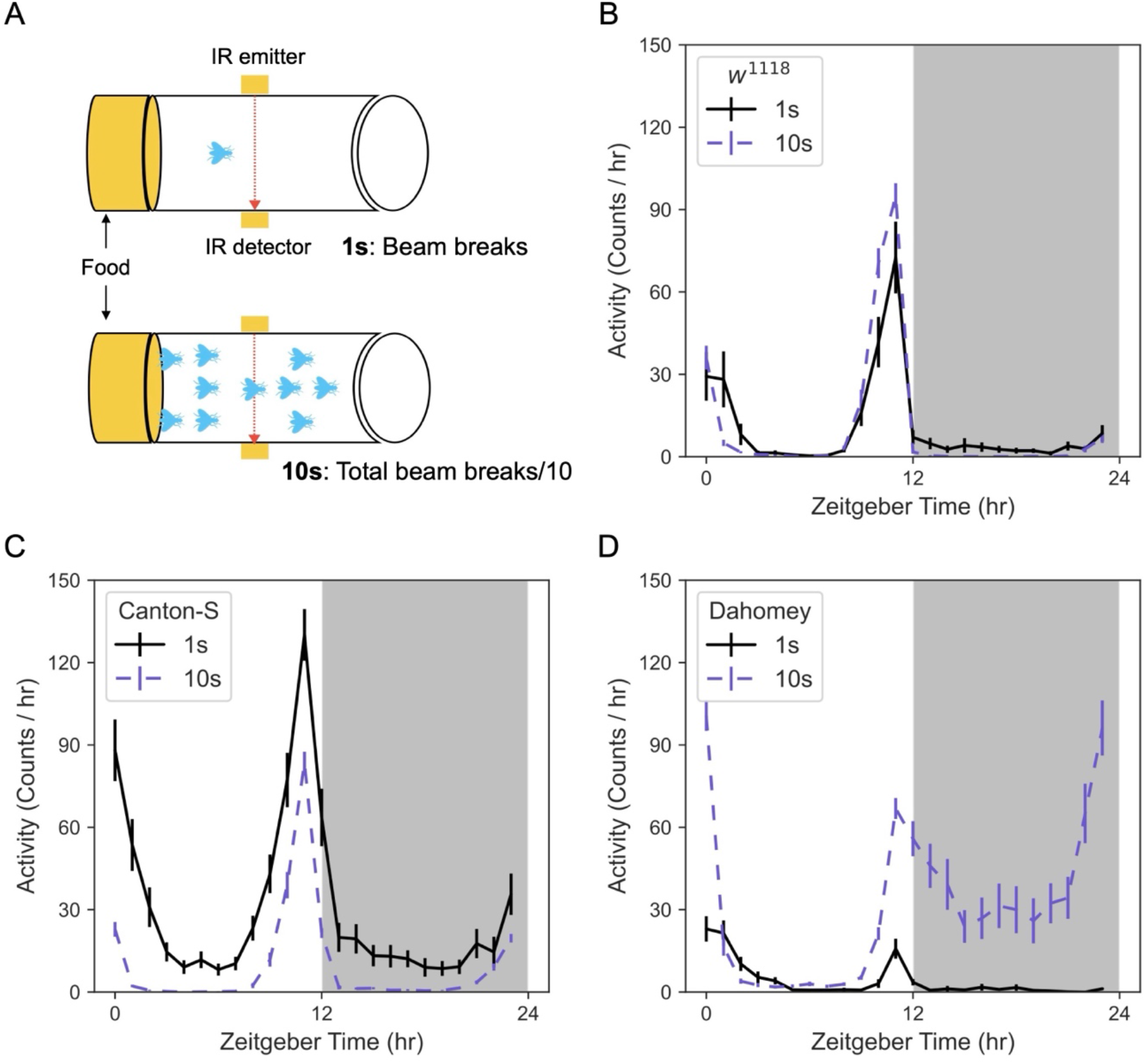
Dahomey flies exhibit hyperactivity in group housing. (A) Schematic of *Drosophila* individual and group locomotor activity measured by the LAM system. Vials containing food and either 1 (1s) or 10 (10s) flies are oriented horizontally and beam breaks are counted by a planar array of infrared (IR) beams that crosses each tube at its midpoint. Activity is reported as the number of beam breaks (counts) per fly. (B-D) Dahomey, but not *w*^*1118*^ or Canton-S, shows a synergistic increase in locomotor activity when housed in groups. Activity over 24 hr of solo- (1s, solid black lines) or group-housed (10s, blue dashed lines) males of the indicated strain: *w^1118^* (B), Canton-S (C), or Dahomey (D). Activity is shown as an average per fly ± s.e.m. in 1 hr bins. N = 21-26 vials per condition.

Since Dahomey males showed robust group hyperactivity, we next asked whether females share the same trait. To rule out an impact from male pheromones that are transferred to the female during mating, we quantified virgin female behavior. Dahomey virgin females behaved similarly to *w^1118^* and Canton-S virgins, showing no signs of group hyperactivity (Fig. S3). These findings suggest that social sensitivity to group housing is a sexually dimorphic phenotype in Dahomey flies.

To understand what happens to Dahomey individuals in group housing, males were monitored by infrared cameras in transparent vials at night. We found that Dahomey males are more active, moving around the food source (Supplemental Video 1). To exclude the possibility that Dahomey males were fighting over a food source, we adjusted the housing conditions to include 2 flies and 2 food sources at either end of a narrow chamber (Fig. S4A). Under these conditions, pair-housed Dahomey males showed a synergistic increase in activity compared to solo-housed flies, while hyperactivity was not observed in pair-housed Canton-S males (Fig. S4B).

Using infrared videography, we observed that the increased activity in paired Dahomey compared with paired Canton-S was closely associated with increased touching behaviors, which included mostly touch-and-run actions along with some tussling and lunging (Fig. S4C, D). Increased chasing behavior was also observed in pair-housed Dahomey (Fig. S4E). The distance between Dahomey males fluctuated frequently due to their increased overall movement (Fig. S4F). These studies support the idea that hyperactivity measured through beam breaks serves as a higher throughput method for estimating social interactions and testing mechanisms that drive social sensitivity.

### Variation in chemosignal sensation, rather than release, mediates group hyperactivity

To determine whether the sensitivity to group housing in Dahomey is due to differences in chemosignal sensation or release, we recorded videos of flies in groups. A test fly was grouped with 9 Canton-S or Dahomey ‘helpers,’ and artificial ‘beam breaks’ were scored manually by counting the number of times that the test fly crossed the midline of the chamber (Fig. 2A). In the positive control, the test Dahomey fly exhibited increased activity when housed with other Dahomey (Fig. 2B). Co-housing with Canton-S helpers also elicited hyperactivity in the Dahomey test fly (Fig. 2C). In contrast, a Canton-S test fly showed decreased activity when co-housed with Dahomey helpers (Fig. 2D), consistent with the prior result observed with group-housed Canton-S (Fig. 1C). Thus, group hyperactivity is due to differences in the response to the presence of other flies, rather than a phenomenon induced by a specific strain. These results suggest that differences in chemosensory reception or signaling, rather than altered chemosignal synthesis or release, between Canton-S and Dahomey are the determining factor for social sensitivity in flies.

**Figure 2:**
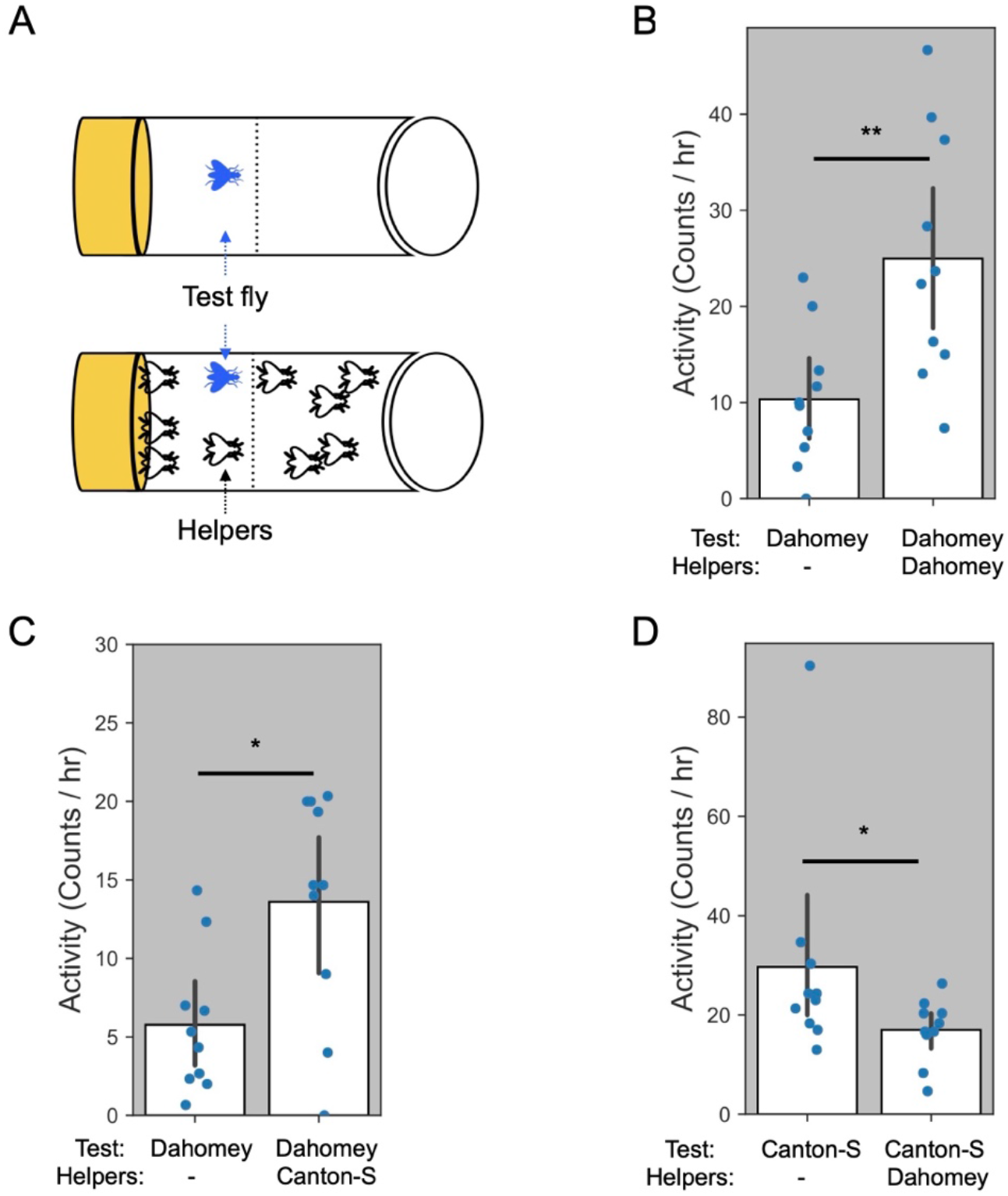
Strain differences in olfaction, rather than in the generation of odors, mediates group hyperactivity. (A) Schematic for testing if fly-specific odors or olfaction contributes to group hyperactivity. A test fly (solid blue) is solo-housed or grouped with 9 helper flies that have clipped wings. (B, C) Dahomey test flies show increased activity when co-housed with either Dahomey (B) or Canton-S (C) helpers. (D) Dahomey helpers do not elicit hyperactivity in a Canton-S test fly. Average per hr ± s.e.m. is shown for a 3-hr test period between ZT 15-18. All test and helper flies were male (N = 10 vials per condition). *, *p* < 0.05; **, *p* < 0.01 (Mann-Whitney U test).

### *Orco* is essential for group hyperactivity in Dahomey

In most insects, including *Drosophila*, antennae are the major sensory organs for olfactory chemosignals^23^. To test whether olfaction is important for Dahomey group hyperactivity, we surgically removed these organs. Antennae-ablated Dahomey did not show group hyperactivity (Fig. 3A, B). Dahomey with surgically removed maxillary palps, a secondary olfactory appendage with 10-fold fewer olfactory sensory neurons (OSNs) than the antennae^24^, continued to exhibit group hyperactivity (Fig. 3C, D), demonstrating that antennae-mediated olfaction is the primary contributor to Dahomey group behavior.

**Figure 3:**
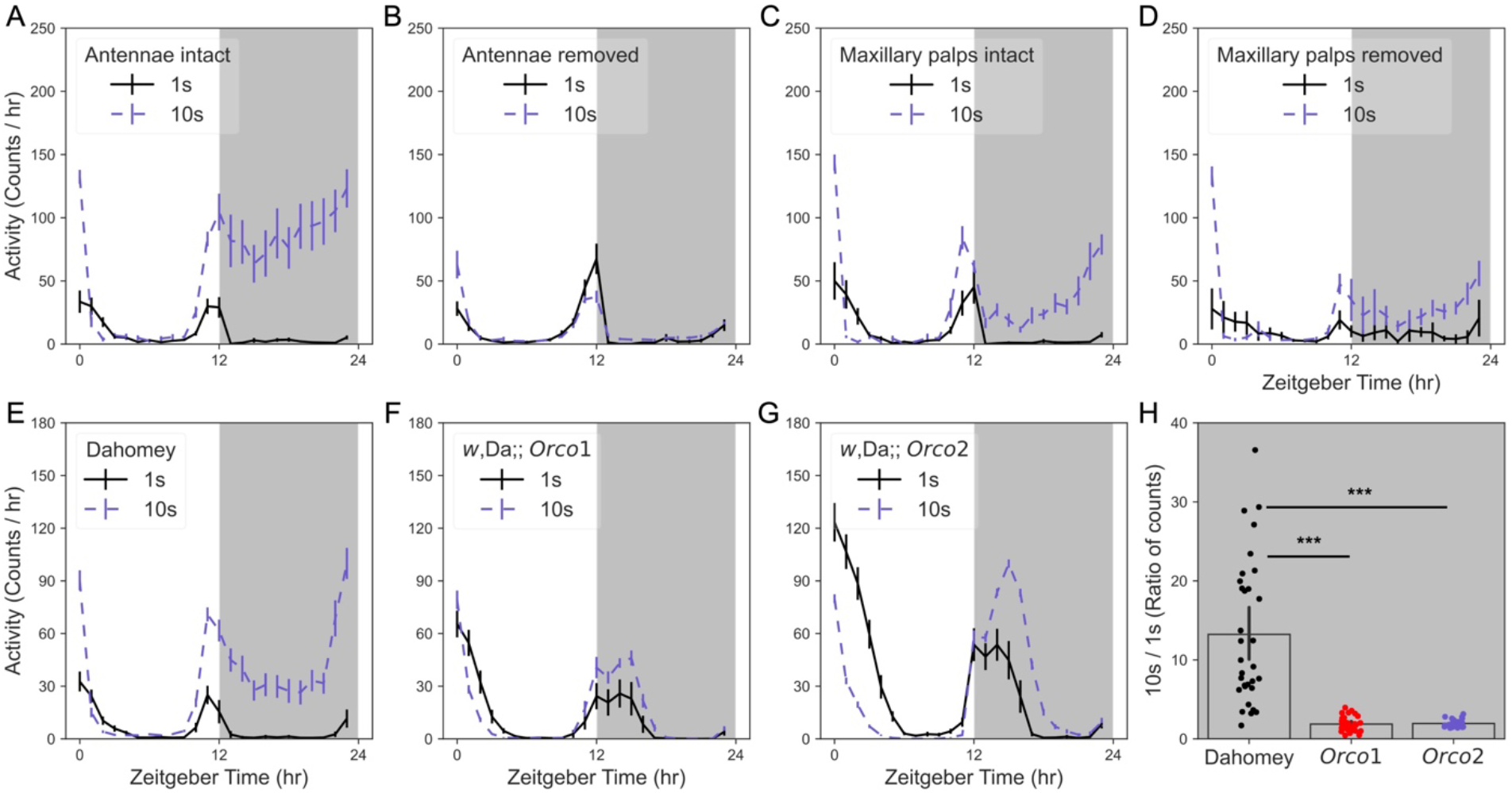
Olfaction is required for group hyperactivity in Dahomey. (A, B) Removal of antennae eliminates group hyperactivity. Activity over 24 hr of Dahomey solo- or group-housed flies with antennae intact (A) or surgically removed (B). (C, D) Removal of maxillary palps does not eliminate group hyperactivity. Activity over 24 hr of Dahomey solo- or group-housed flies with maxillary palps intact (C) or surgically removed (D). Mock-treated controls are shown in A and C. (E-G) *Orco* mutants show reduced sensitivity to group housing. Activity over 24 hr of *Orco1* (F) and *Orco2* (G) mutants in a *w*Dahomey background. Since the *Orco* mutants reintroduce the *w* gene, red-eyed Dahomey is used as the control (E). Activity is shown as an average per fly ± s.e.m. in 1 hr bins. (H) Response to group-housing in different genotypes, expressed as a ratio of activity per fly in groups vs. solo-housed from the 12-hr dark period. Males used in all experiments. N = 16-20 vials per condition (A, B), 7 vials per condition (C, D) or 32 vials per condition (E-G). ***, *p* < 0.001 (Dunn’s test with Bonferroni correction). *w*, Da = *w*Dahomey background.

OSNs include neurons that express odorant receptors (ORs) and ionotropic receptors that are used for specific odor detection^25^. In particular, the broadly expressed ORCO (also known as OR83b) is highly conserved among insect species, and chaperones other ORs^26^. To further validate that olfaction is necessary for group hyperactivity, *Orco* mutations (*i*.*e*., *Orco1* and *Orco2*) were introduced into the *w*Dahomey (Dahomey bearing *white* gene mutation) background. *Orco* mutations significantly reduced the group-housing effects in Dahomey (Fig. 3E-G). To account for differences in baseline behavior of solo-housed flies from different genotypes, the effect of group-housing on activity is expressed as a ratio of the activity per fly observed in groups, divided by the activity quantified in singly housed animals (10s/1s), typically from the 12-hour dark period (Fig. 3H). Since the *Orco1* mutant had a lower baseline group activity (Fig. 3F), we subsequently used *Orco1* in genetic rescue studies.

Given that group hyperactivity was effectively inhibited in *Orco1*, we next tested if the phenotype could be restored by genetically rescuing *Orco* expression. The first approach we used was re-expressing exogenous *Orco* in *Orco1* flies using the UAS/GAL4 binary system^26^. Compared to Dahomey, negative controls harboring the *Orco*1 mutation in the Dahomey background, with or without one of the UAS/GAL4 transgenic elements, did not exhibit robust group hyperactivity (Fig. 4A-D, F). *Orco*-rescued flies displayed ∼6-fold greater nighttime activity in the group-housed condition compared to being housed individually, demonstrating a partial rescue of the phenotype when comparing with Dahomey and the non-rescue controls (Fig. 4E, F).

**Figure 4:**
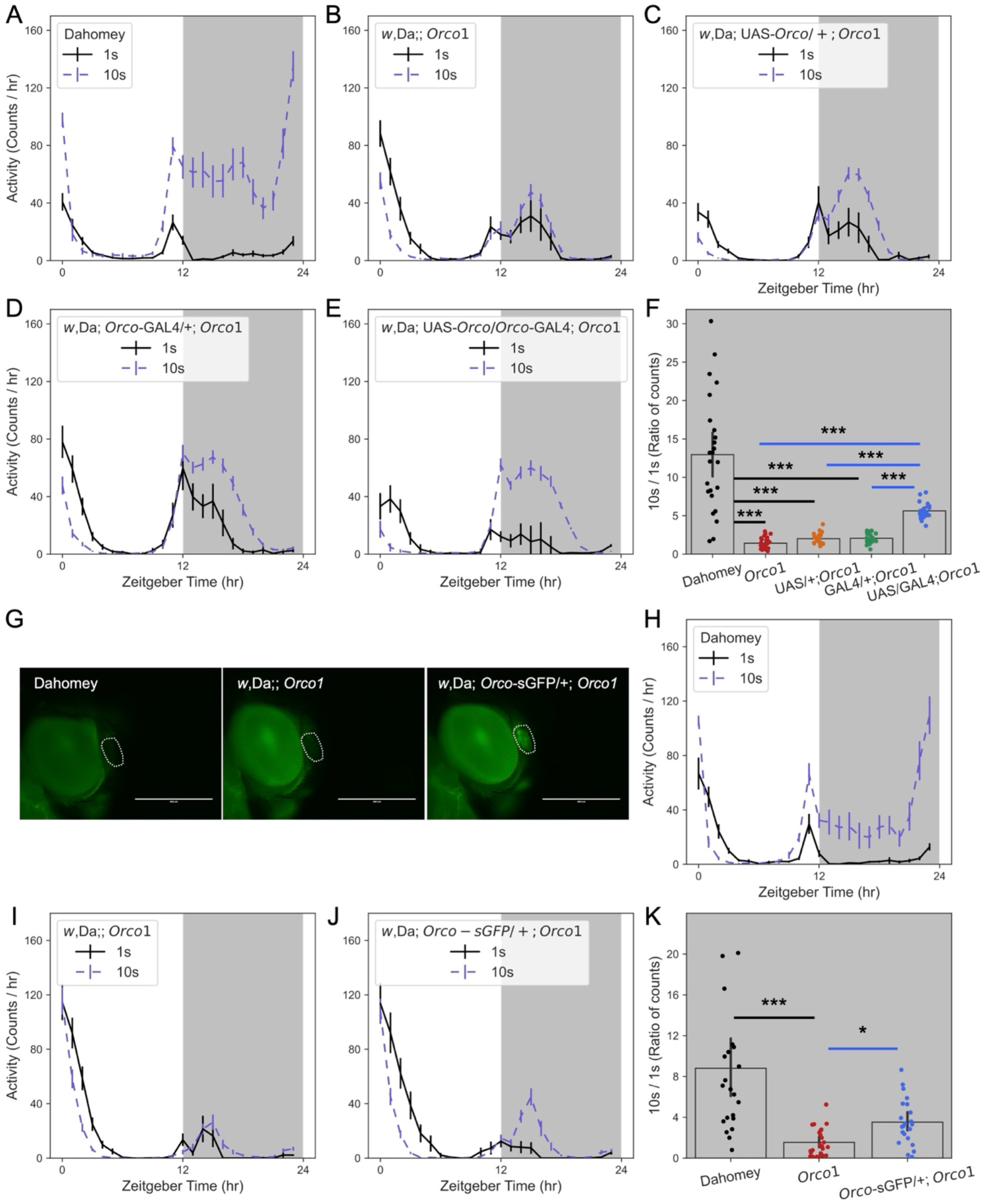
Rescue of group hyperactivity in *Orco* mutants. (A-E) Activity over 24 hr of Dahomey (A), the *Orco1* mutant (B), UAS*-Orco/+; Orco1* (C), *Orco-* GAL4/*+; Orco1* (D), or *Orco*-rescued flies (E). (F) Response to group-housing in different genotypes, expressed as a ratio of activity per fly in groups vs. solo-housed from the 12-hr dark period. (G) Orco-sGFP expression in the fly antenna (circled). Scale bars: 400 μm. (H-J) Activity over 24 hr of Dahomey (H), the *Orco1* mutant (I), or *Orco-sGFP/+*; *Orco1* (J). (K) Response to group-housing in different genotypes, expressed as a ratio of activity per fly in groups vs. solo-housed from the 12-hr dark period. Males used in all experiments. For 24-hr actograms, activity is shown as an average per fly ± s.e.m. in 1 hr bins. N = 19-25 vials per condition. *, *p* < 0.05; ***, *p* < 0.001 (Dunn’s test with Bonferroni correction). *w*, Da = *w*Dahomey background.

Rescue of the *Orco1* mutant with an *Orco* transgene also partially restored group hyperactivity. The *Orco-sGFP* construct contains an intact *Orco* gene sequence tagged with superfold-GFP^27^, which shows the potential for expressing and localizing *Orco* in antennae. GFP signal was observed in *Orco1* antennae (Fig. 4G), indicating that the *Orco-sGFP* insertion was successfully expressed. Group hyperactivity could be partially restored in *Orco1* with ORCO-sGFP expression (Fig. 4H-K). However, the rescue effect was not as strong as the results with the UAS/GAL4 method, possibly because a single copy of *Orco* is not sufficient to restore intact olfactory function. Homozygous *Orco-sGFP* is lethal and could not be tested.

### *Orco*^*+*^ and *Or43a*^*+*^ neurons contribute to group hyperactivity in Dahomey

In addition to the broadly expressed *Orco* in antennae, odorant receptors (ORs) are expressed in specific *Orco*^*+*^ neurons^28^. In *Drosophila*, the detection of odorants depends on heterodimers consisting of ubiquitously expressed Orco and an ORN-specific OR^29^. ORNs project to the central brain to form different glomeruli, establishing connections between odorant signals and the brain^28^. To verify if *Orco*^*+*^ neurons are necessary for group hyperactivity, we silenced *Orco*^*+*^ neurons in a Dahomey background (Fig. 5A). Tetanus toxin (TNT) cleaves Synaptobrevin (Syb) in synaptic vesicles, leading to the blockade of neurotransmission^30^. Expressing *TNT* in *Orco*^*+*^ neurons eliminated group hyperactivity in Dahomey (Fig. 5B-D). Expression of *Kir2*.*1* induces hyperpolarization of *Orco* neurons, leading to similar results (Fig. 5E-H), suggesting that *Orco*^*+*^ neurons gatekeep chemosignal-induced group hyperactivity. Expressing the proapoptotic gene *Reaper* also effectively inhibited group hyperactivity by promoting cell death of *Orco*^*+*^ neurons (Fig. 5I-L).

**Figure 5:**
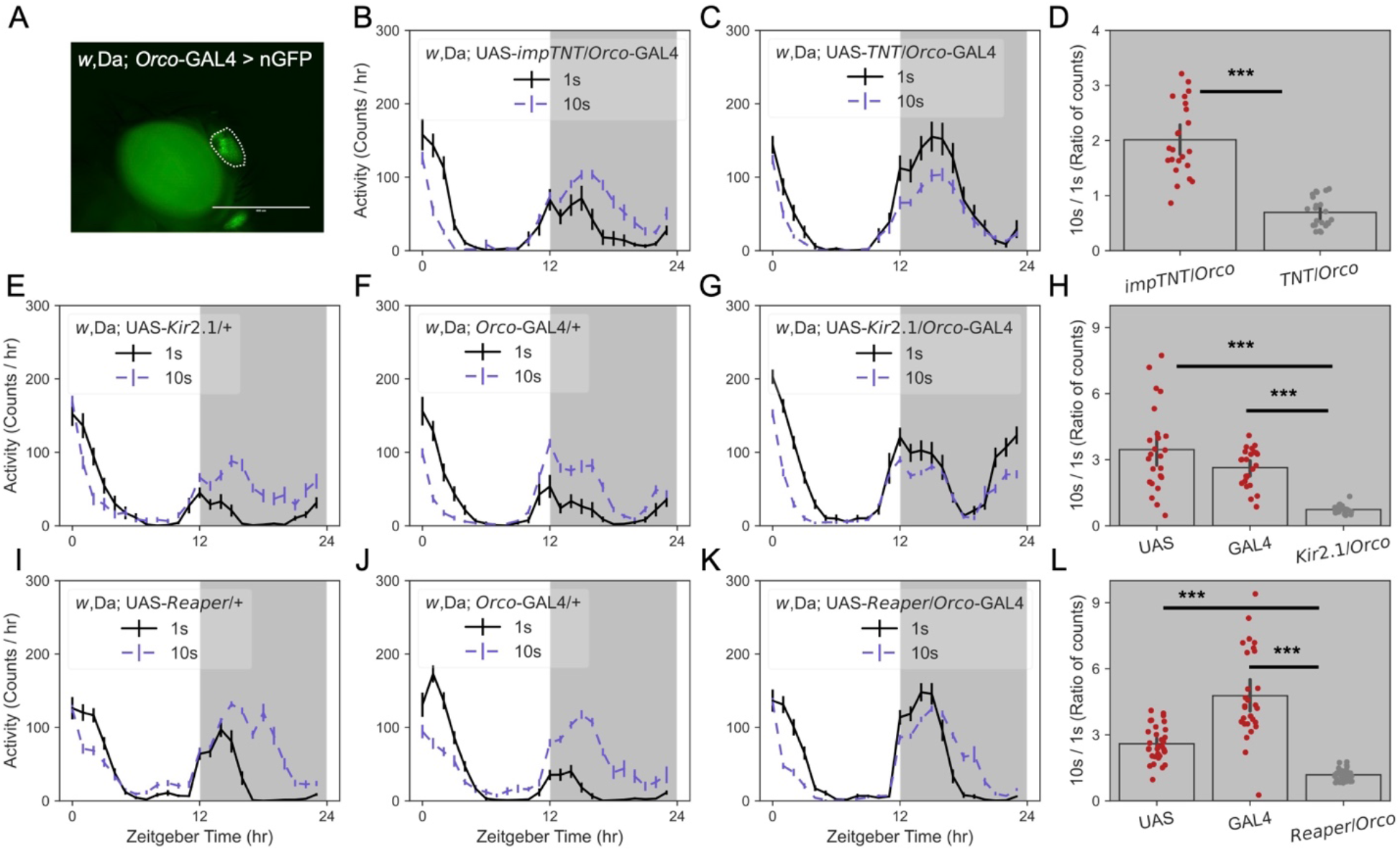
Silencing of *Orco* neurons inhibits group hyperactivity in Dahomey. (A) *Orco-*GAL4 drives nGFP expression in antennae. Scale bar: 400 μm. (B-D) Blocking neurotransmission in *Orco*^*+*^ neurons by expressing TNT (C) eliminates group hyperactivity (D). Expression of an inactive form (impTNT) is used as a control (B). (E-H) Silencing *Orco* neurons by expressing Kir2.1 (G) inhibits group hyperactivity (H). Controls harbor only the *Kir2*.*1* (E) or the *GAL4* (F) transgene. (I-L) Inducing apoptosis of *Orco* cells by expressing Reaper (K) represses group hyperactivity (L). Controls harbor only the *Reaper* (I) or the *GAL4* (J) transgene. Males used in all experiments. For 24-hr actograms, activity is shown as an average per fly ± s.e.m. in 1 hr bins. N = 22-24 (B-H) or 28-35 (I-L) vials per condition. ***, *p* < 0.001 (Dunn’s test with Bonferroni correction or Mann-Whitney U test). *w*, Da = *w*Dahomey background.

To identify specific *Or-X*^*+*^ neurons that affect group hyperactivity, we screened 20 different ORN drivers in a Dahomey background using *Reaper, Kir2*.*1*, and *TNT* (Supplemental Table 1). Inhibition of *Or43a*^*+*^ neurons consistently decreased group hyperactivity (Fig. 6 & Supplemental Table 1). To test if olfactory signaling through *Or43a*^*+*^ neurons is sufficient to mediate group hyperactivity, we expressed Orco in *Or43a*^*+*^ neurons of *Orco*1 mutants using the UAS/GAL4 system. Restoring Orco only in *Or43a*^*+*^ neurons did not rescue group hyperactivity in the *Orco1* mutant (Fig. S5), implying that additional ORNs are required.

**Figure 6:**
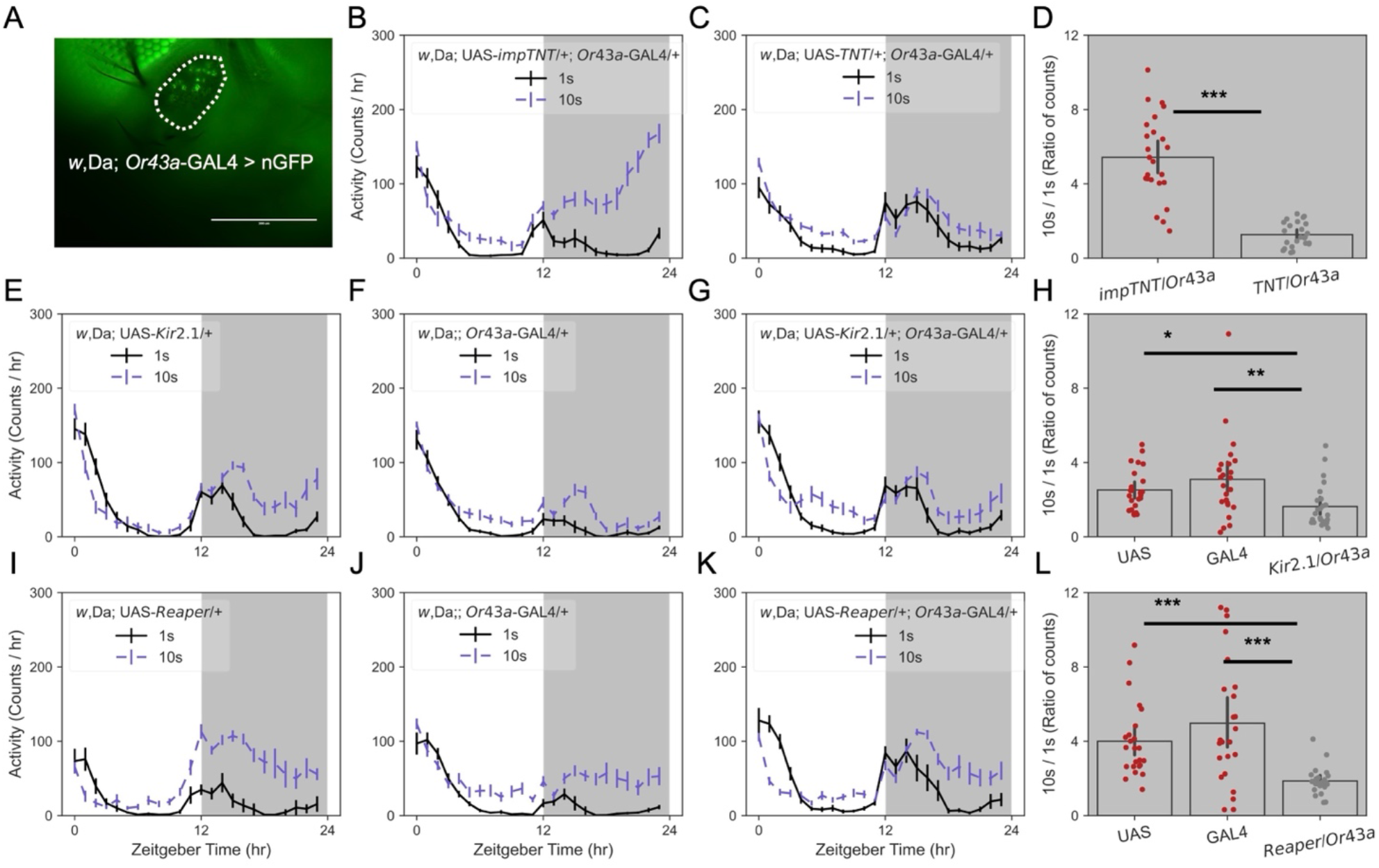
Silencing of *Or43a*^*+*^ neurons decreases group hyperactivity in Dahomey flies. (A) *Or43a-*GAL4 drives nGFP expression in antennae. Scale bar: 400 μm. (B-D) Blocking neurotransmission in *Or43a*^*+*^ neurons by expressing TNT (C) represses group hyperactivity (D). Expression of an inactive form (impTNT) is used as a control (B). (E-H) Silencing *Or43a*^*+*^ neurons by expressing Kir2.1 (G) inhibits group hyperactivity (H). Controls harbor only the *Kir2*.*1* (E) or the *GAL4* (F) transgene. (I-L) Inducing apoptosis of *Or43a*^*+*^ cells by expressing Reaper (K) decreases group hyperactivity (L). Controls harbor only the *Reaper* (I) or the *GAL4* (J) transgene. For all studies, N = 23-24 vials per condition. Males used in all experiments. For 24-hr actograms, activity is shown as an average per fly ± s.e.m. in 1 hr bins. N = 23-24 vials per condition. *, *p* < 0.05; **, *p* < 0.01; ***, *p* < 0.001 (Dunn’s test with Bonferroni correction or Mann-Whitney U test). *w*, Da = *w*Dahomey background.

### Dahomey group hyperactivity is not induced by canonical odors

Given the prominent role of olfaction in Dahomey group hyperactivity, it is possible that volatile chemosignals drive social behavior. Coincidentally, cVA is an important volatile pheromone released by male flies that can be perceived at a long distance^31^. The characteristics of cVA in promoting male-male aggression have been reported^32^. However, the physical separation of a single fly from a stimulus source (10 flies) without blocking airflow (Fig. S6A) did not induce robust hyperactivity (Fig. S6B). This implies that the olfactory cues from group-housed Dahomey might function within a short range between flies, or that additional cues are necessary to drive activity. In contrast to cVA, CH compounds are deposited as a thin layer on the cuticle, and only a few are known to be slightly volatile^33^. To investigate whether CHs induce group hyperactivity, we genetically disrupted the synthesis of CHs *in vivo*. The *Desalt1* driver expresses GAL4 in both oenocytes and the male ejaculatory bulb (Fig. S7A), in which CHs and cVA are synthesized, respectively. Ablating larval- or pupal-stage oenocytes arrested development and caused lethality. Expressing *Reaper* or *Hid* in adults can promote the apoptosis of oenocytes without affecting cVA synthesis^14,34^. Thus, the temperature-sensitive GAL4 inhibitor, Tub-GAL80^ts^, was introduced to bypass developmental lethality (Fig. S7B). Dahomey flies with ablated adult oenocytes still exhibited group hyperactivity (Fig. S7C-E), suggesting that group hyperactivity is not mediated by CHs.

### Genetic variation in Dahomey *Or43a* may affect group hyperactivity

The cumulative results support the idea that signaling through *Or43a*^*+*^ neurons is necessary but not sufficient for robust Dahomey group hyperactivity. We next tested whether the OR43a receptor mediates the phenotype by testing CRIMIC insertions of *Or43a*, where the SA-T2-GAL4-Poly(A) cassette arrests *Or43a* transcription^35^. Through 10 generations of outcrossing, the genetic background of *Or43a*^CR60034^ was replaced with *w*Dahomey. *Or43a*^CR60034^ successfully drove RedStinger expression in antennal neurons (Fig. 7A), implying that the cassette functions as intended. Similar to the inhibition of *Orco*^*+*^ or *Or43a*^*+*^ neuronal signaling, the CR60034 insertion increased the baseline activity of solo-housed flies (Fig. 7B, C). Compared to *w*Dahomey, *Or43a* loss-of-function significantly decreased the synergistic effect of group-housing on activity (Fig. 7D), suggesting that the receptor at least partially contributes to the phenotype.

**Figure 7:**
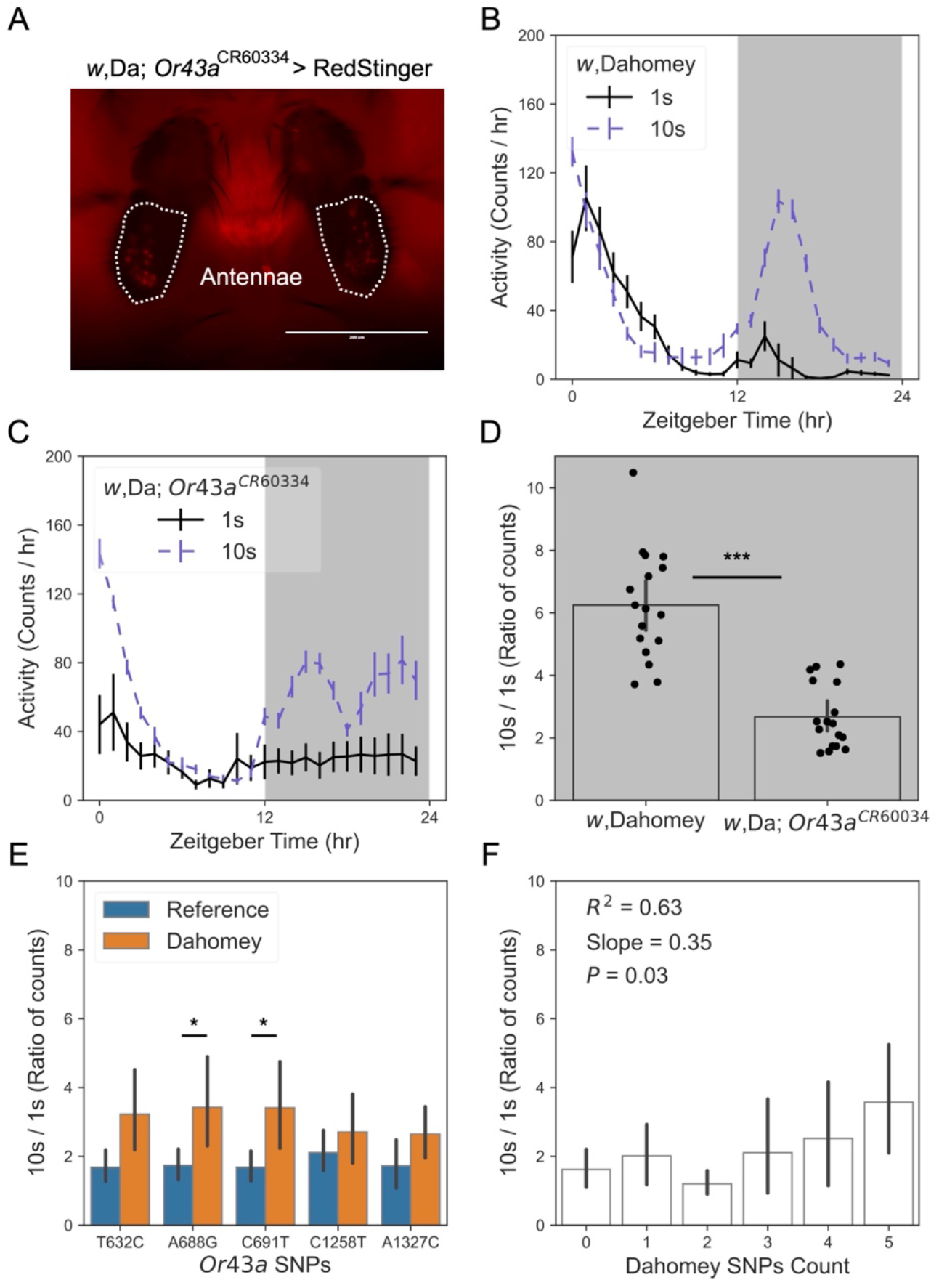
*Or43a* is involved in the regulation of group activity in flies. (A) The SA-T2-GAL4-Poly(A) cassette in an intron of *Or43a* drives RedStinger expression in antennae. Scale bar: 200 μm. (B-D) The *Or43a* insertion mutant (C) displays reduced group hyperactivity (D) compared to the *w*Dahomey control (B). For 24-hr actograms, activity is shown as an average per fly ± s.e.m. in 1 hr bins. N = 17 vials per condition. (E) Effect of Canton-S vs. Dahomey SNPs on nighttime group hyperactivity in 135 lines from the DGRP collection. Transcriptional start site of *Or43a*: +1. (F) The linear regression of nighttime group hyperactivity of the DGRP lines, categorized based on the number of Dahomey *Or43a* SNPs that each line contains. *R*^2^: Correlation coefficient; *p* = 0.03 indicates the rejection of the null hypothesis that the slope ≤ 0. Males used in all experiments. *, *p* < 0.05; ***, *p* < 0.001 (Mann-Whitney U test). *w*, Da = *w*Dahomey background.

Since the *Orco* and *Or43a* genes are important for regulating group hyperactivity, we sequenced these two gene regions in Canton-S and Dahomey. *Orco* was found to be highly conserved, with identical sequences in both strains. However, 11 single nucleotide polymorphisms (SNPs) were identical in *Or43a* between Canton-S and Dahomey. To ask whether genetic variation in the *Or43a* gene might contribute to group hyperactivity, we examined the *Drosophila* Genetics Reference Panel (DGRP), a collection of ∼200 inbred fly strains used for GWAS^36,37^. Interestingly, 5 out of the 11 SNPs between Canton-S and Dahomey are well represented in the DGRP. We quantified activity in solo- and group-housed flies from 135 lines in the DGRP collection (Supplemental Table 2). By grouping the results based on the *Or43a* SNPs present, we found that 2 of the 5 SNPs in *Or43a* are significantly associated with group hyperactivity, with all 5 SNPs individually trending toward increased group hyperactivity when the Dahomey SNP was present (Fig. 7E). We also categorized the DGRP results based on the number of Dahomey *Or43a* SNPs present, from 0 (most CS-like) to 5 (most Dahomey-like). These results showed a significant correlation between the number of Dahomey-containing *Or43a* SNPs present and the level of group hyperactivity (Fig. 7F), suggesting that natural genetic variation within this gene contributes to behavioral responses to social environment.

## Discussion

Dahomey males exhibit robust group hyperactivity, contrasting with other tested *D. melanogaster* strains. Video analyses suggest that the increased activity is due to social behaviors including touching and chasing. The interactions did not appear to be male-male courtship since canonical courtship behaviors such as wing vibrations were not observed. However, future work might better define the specific social interactions that are occurring among male Dahomey through higher resolution videography and machine learning methods for recognizing patterns of social behavior^38^.

Our results suggest that differences in olfactory chemosensation, rather than the generation of chemosignals, underlie the divergent responses to group housing in Dahomey males. Sexually dimorphic olfaction might also play a role in the sex dependence of social sensitivity in Dahomey. The *fruitless* gene is responsible for the development of many brain sex differences and a mosaic analysis of *fruitless*^*+*^ neurons previously revealed that dimorphic clones overlap mainly in olfactory neurons over other sensory neurons^39^. Additionally, the male-specific growth of some *Drosophila* glomeruli within the olfactory bulb can be prevented by the ectopic expression of female-type *transformer* in males, demonstrating glomerular sexual dimorphisms^40^.

Although *Or43a* contributes to social sensitivity, other ORs might also be needed since ORCO re-expression in *Or43a*^*+*^ neurons in an *Orco* mutant background was not sufficient to rescue Dahomey group hyperactivity. Our studies did not identify the specific olfactory chemosignals that are important for driving Dahomey male social interactions, but a previous study showed that OR43a recognizes cyclic molecules with a polar group, including benzylaldehyde, benzyl alcohol, cyclohexanol, and cyclohexanone^41^. Future studies might examine whether these ligands affect locomotor or social behaviors.

Our behavioral analyses predominantly rely on comparing the response of an individual fly to group housing by expressing it as a ratio of per fly activity in groups to solo-housed activity, within genotype. However, we also observed some consistent between-genotype differences in behavior. Inhibition of olfaction—by using *Orco* or *Or43a* mutants, silencing of *Orco*^*+*^ or *Or43a*^*+*^ neurons, or surgically removing antennae—increases solo-housed activity. These results might reflect previous studies showing that chronic isolation reduces individual sleep^6^. In ants, knockout of *Orco* impairs the fitness of individuals within groups^42^. It is possible that inhibition of *Orco* signaling simulates chronic isolation to alter behavior. Increased solo-housed activity was not evident with disruption of CH synthesis (Fig. S7), possibly due to social experience before the genetic ablation of oenocytes was induced.

Although the genetic basis underlying strain-specific differences in group behaviors is not well understood, it is possible that variations in specific ORs determine behavioral responses to the social environment. One Dahomey *Or43a* SNP results in a V254A substitution in the receptor that is not found in Canton-S, but is commonly observed in the OR43a protein sequence of other *Drosophila* species. Various species of *Drosophila* can show differences in coding of the same odors^43^. V254A was not present in the DGRP, but a comparative genomic analysis of the olfactory genes between Dahomey and other strains might reveal key genetic features that modulate social behavior.

Although it is unclear whether Dahomey *Or43a* SNPs are functional or causal for behavioral phenotypes, our studies using the DGRP collection suggest that lines containing more ‘Dahomey-like’ *Or43a* sequences show greater group hyperactivity. Interestingly, when examining the effect of each of the 26 *Or43a* SNPs in the DGRP, only 2 SNPs show a statistically significant effect—and they are both present in Dahomey. Although these two SNPs are in exons, they result in silent mutations. Nonetheless, synonymous mutations can impact phenotypes through other processes^44^. Future work could test whether specific SNPs, perhaps generated by CRISPR, is sufficient to induce group hyperactivity in a Canton-S background or repress the phenotype in a Dahomey background.

## Acknowledgments

We thank Dr. Scarlet Park for helpful comments on this manuscript. This work was funded by the NIH (R01DC020031, W.W.J.).

## Author Contributions

Conceptualization, Methodology, Data collection, and Data Analysis: B.W., E.S.K., and W.W.J.; Supervision: W.W.J.; Writing—original draft: B.W.; Writing—review and editing: B.W. and E.S.K.

## Materials & methods

### *Drosophila* strains and rearing

*Orco*1 (#23129), *Orco*2 (#23130), UAS-*Orco* (#23145), *Orco*-GAL4 (#26818), CRIMIC*_Or43a_*TG4.1 (#94381), *Or43a*-GAL4 (#9974), *Desat1*-GAL4 (#65405), and the DGRP lines were obtained from the Bloomington *Drosophila* Stock Center. *Orco-sGFP* (v318654) was from the Vienna *Drosophila* Resource Center. All lines have been backcrossed into the indicated line (*w*Dahomey or *w*Canton-S) for at least 10 generations. Flies were reared at 25 °C in a 12-/12-hour light/dark cycle. Flies with *tub-GAL80*^*ts*^ insertions were raised at 18 °C during larval and pupal development to inhibit GAL4 expression. Flies were typically collected up to 2 days post-eclosion, sorted under light CO_2_ anesthesia, and maintained as single sex populations until used for behavioral studies. Virgin females were collected within 4-6 hr of eclosion.

### Activity measurement

The Locomotor Activity Monitor LAM25H (TriKinetics, Princeton, MA) was used to measure the activity of vial-housed flies. Flies (∼7-days post-eclosion) were randomly allocated into the indicated condition (typically 1 or 10 flies per vial). Activity data were recorded as the number of beam breaks over time. Results from the group-housed condition were analyzed as the beam break counts per fly. Fly responses to group housing were expressed as a ratio of the activity per fly in groups divided by the average activity when singly housed.

### Videography

Flies were acclimated in transparent vials or arenas for several hours prior to lights off, and then recorded by infrared cameras, typically for 3 hr starting at ZT 14 or 15. Supplemental Video 1 shows 30 min of original video sped up by 32×. For results in Figure 2, the wings of helper cohorts were clipped, to facilitate manual scoring of activity of the test fly.

### Statistical analysis

*P* values among multiple comparisons were determined by Dunn’s test with Bonferroni correction following one-way ANOVA (Kruskal-Wallis H-test), using the Scikit-posthocs package in Python. The *P* values between two groups were determined by Mann-Whitney U test; the linear regression was conducted by using SciPy.Stats package in Python. The 24-hour activity curves and bar graphs were made with Python, and data shown are typically mean ± s.e.m.

