## Supplemental Figures for "Chemosensation drives divergent social behavior in *Drosophila*"

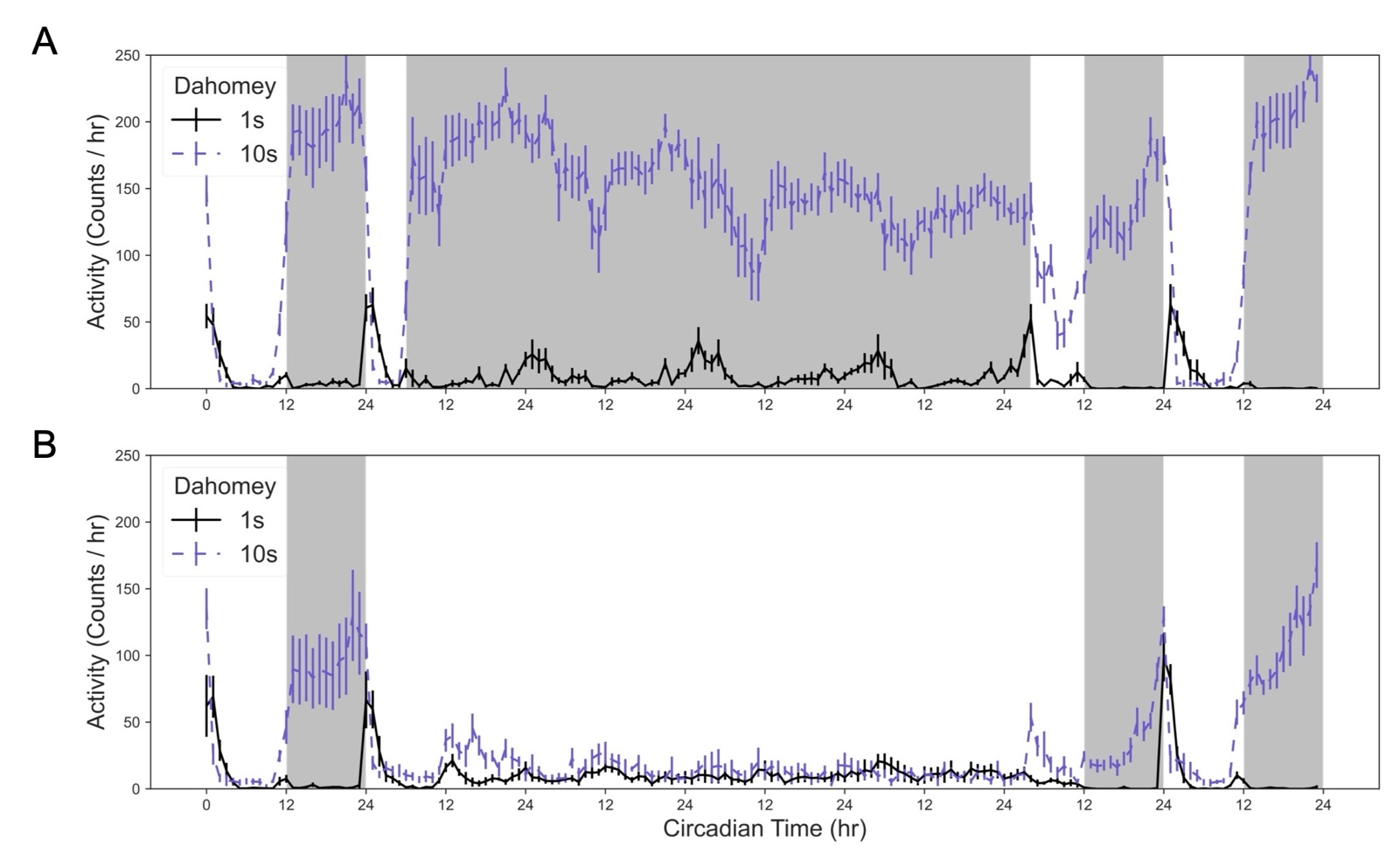

**Figure S1: Light inhibits Dahomey group hyperactivity.**

(A) Dahomey males showed continuous group hyperactivity during >3 days of constant darkness. (B) Group hyperactivity was suppressed by light. Activity is shown as an average per fly ± s.e.m. in 1 hr bins. N = 6-8 vials of Dahomey males per condition.

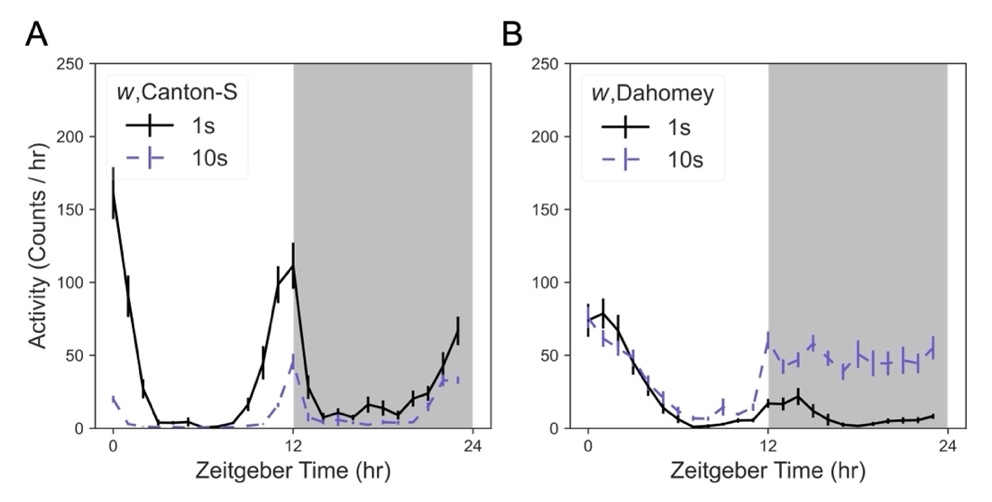

**Figure S2: The *white* mutation does not affect Dahomey group hyperactivity.**

(A, B) Similar to the parental Canton-S and Dahomey strains, Canton-S males with the *white* mutation (A) displayed decreased activity when they were group-housed, while Dahomey with the *white* mutation exhibited nighttime group hyperactivity (B). Activity is shown as an average per fly ± s.e.m. in 1 hr bins. N = 29-32 vials per condition.

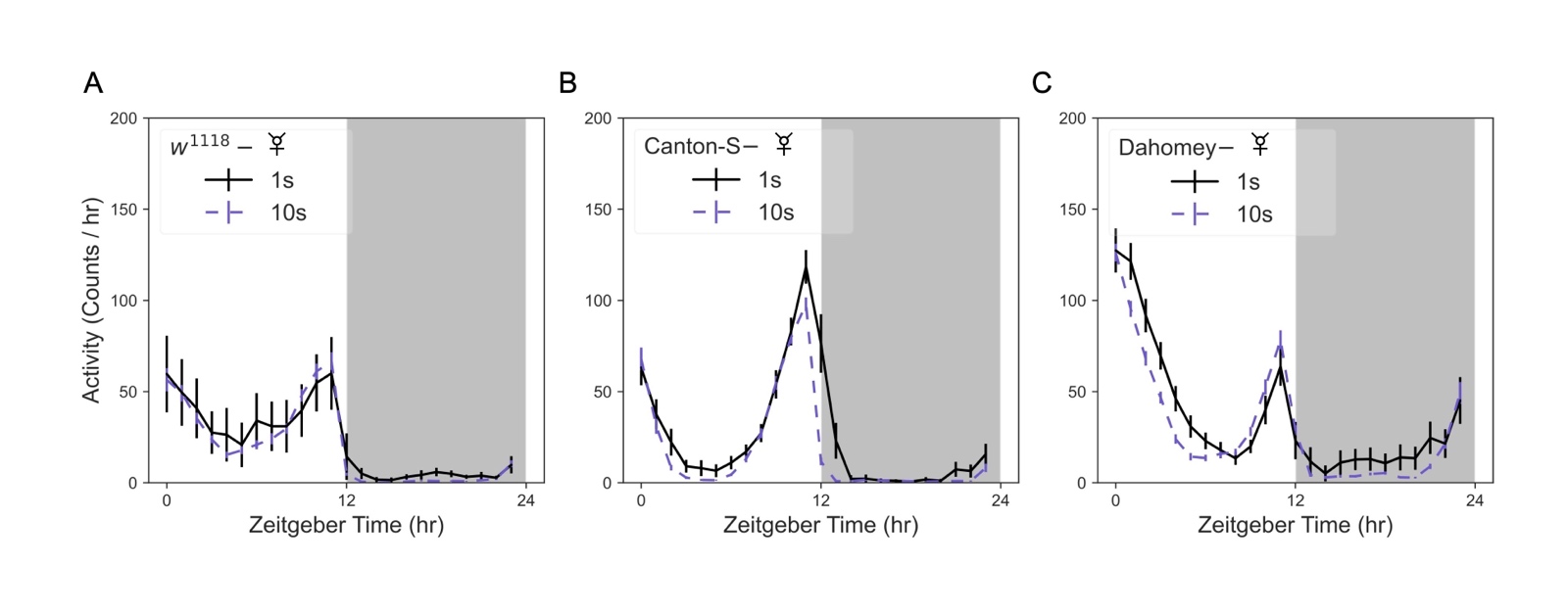

**Figure S3: Females do not show group hyperactivity.**

(A-C) Activity over 24 hr of *w*^1118^ (A), Canton-S (B), and Dahomey (C) virgin females did not show any group hyperactivity. Activity is shown as an average per fly ± s.e.m. in 1 hr bins. N = 18-23 vials per condition.

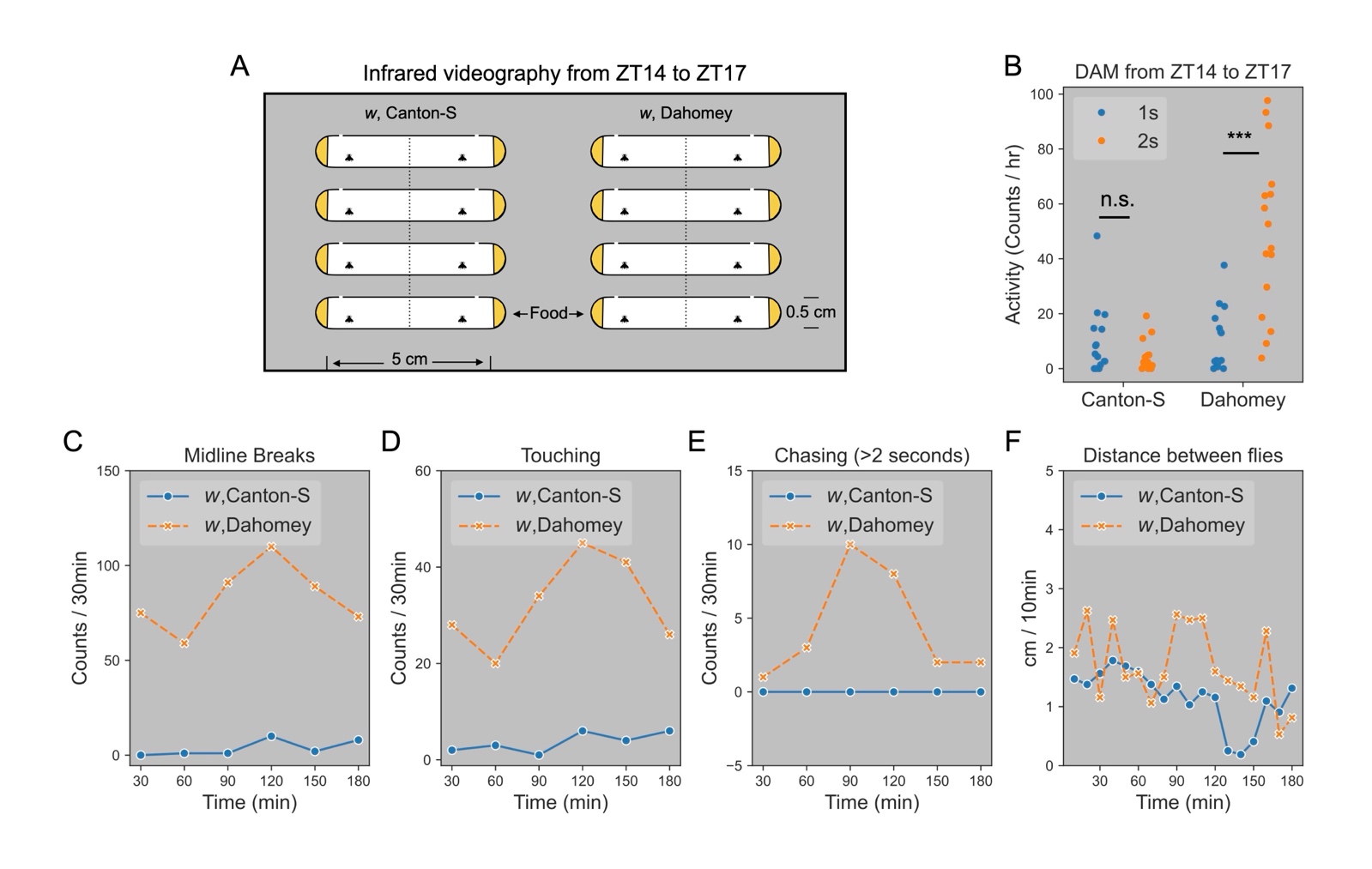

**Figure S4: Infrared videography for group-housed flies.**

(A) Flies were housed in 5 cm × 0.5 cm tubes, providing food and air holes on both sides. After 1 day of acclimation, the infrared camera was set up to record 3-hour video from ZT14 to ZT17. (B) DAM system validates that group housing of 2 Dahomey males still induces hyperactivity. For infrared videography, the midline breaks (C), touching (D), and chasing (E) behaviors were counted per 30 minutes. Distance between flies (F) was measured per 10 minutes. ***, *p* < 0.001 (Mann-Whitney U test).

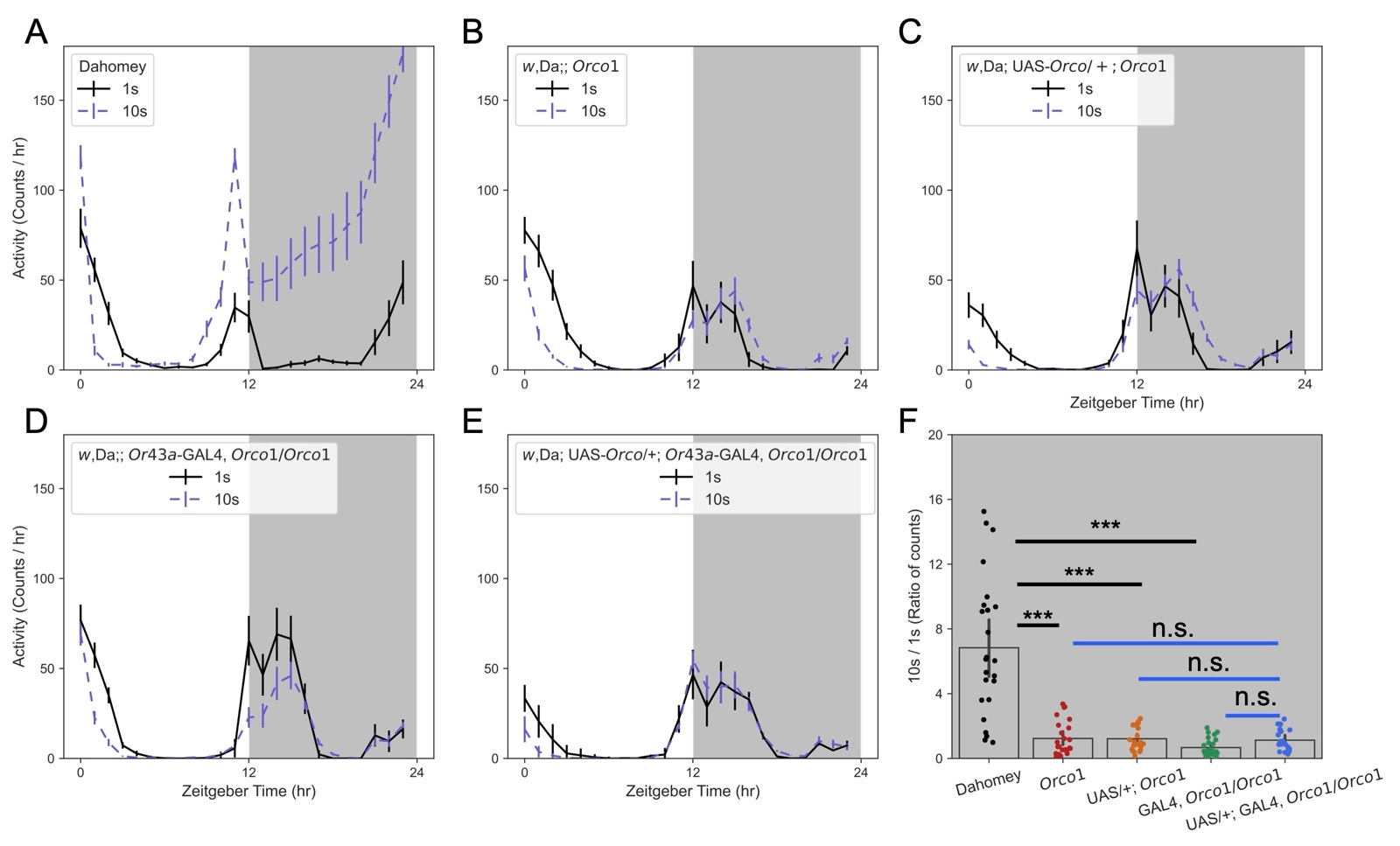

**Figure S5: Expressing *Orco* in *Or43a^+^* neurons does not rescue group hyperactivity in *Orco*1 mutants.**

Activity over 24 hr locomotor activity was measured in Dahomey (A), *Orco1* (B), UAS*-Orco/+;* *Orco1*(C), *Or43a-*GAL4*, Orco1/Orco1* (D), and *Orco*-rescued flies (E). Activity is shown as an average per fly ± s.e.m. in 1 hr bins. (F) Response to group-housing in different genotypes, expressed as a ratio of activity per fly in groups vs. solo-housed from the 12-hr dark period. Males used in all experiments. N = 21-24 vials per condition. ***, *p* < 0.001 (Dunn’s test with Bonferroni correction).

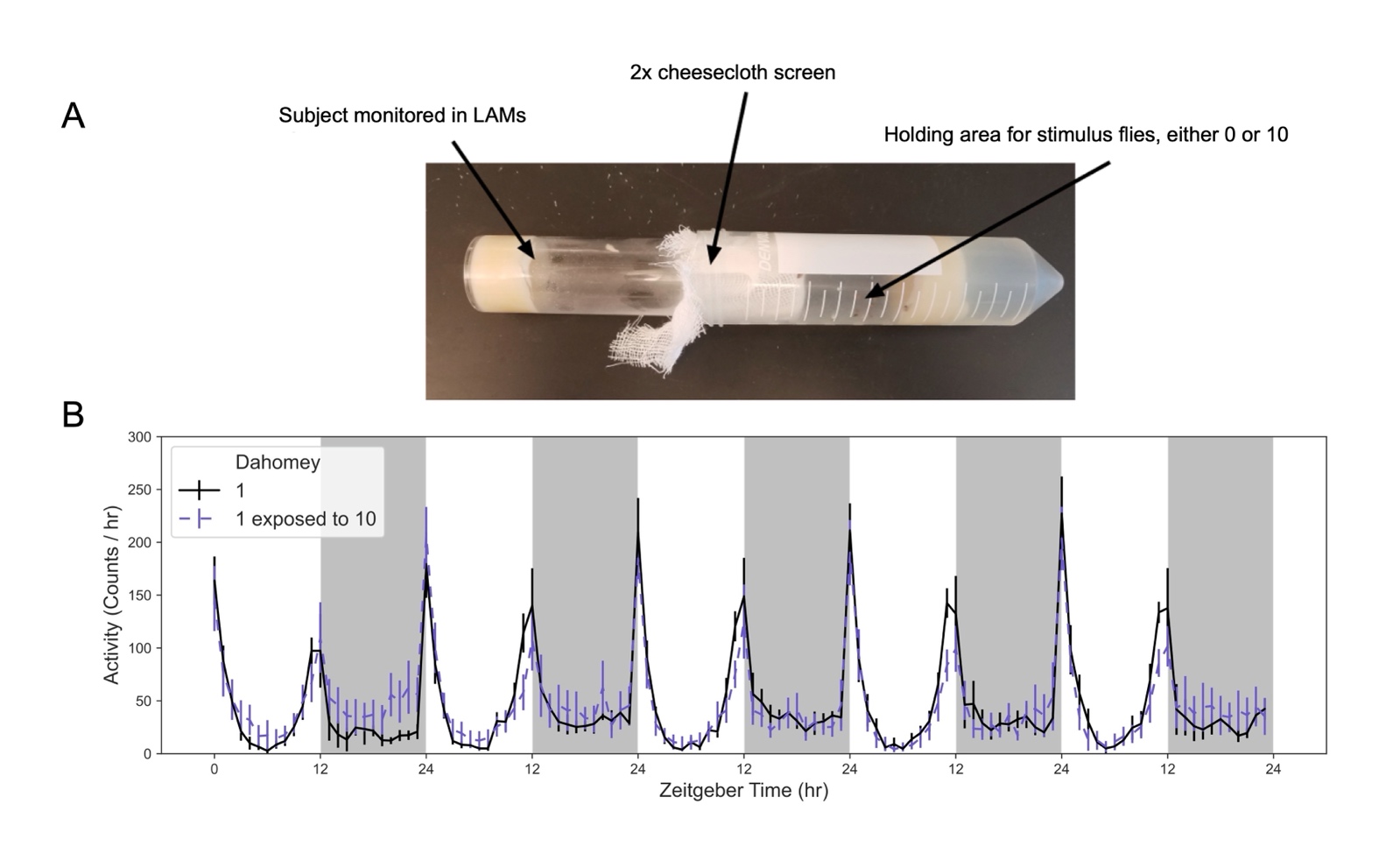

**Figure S6: Group hyperactivity is not induced by long-range odors**

(A) A specialized vial for physically separating a test fly from other animals, while sharing potential long-range odors. (B) No obvious difference was observed between test flies housed with or without other flies. Activity of the test fly was quantified in the LAM system. N = 6 vials for single and group-housed males.

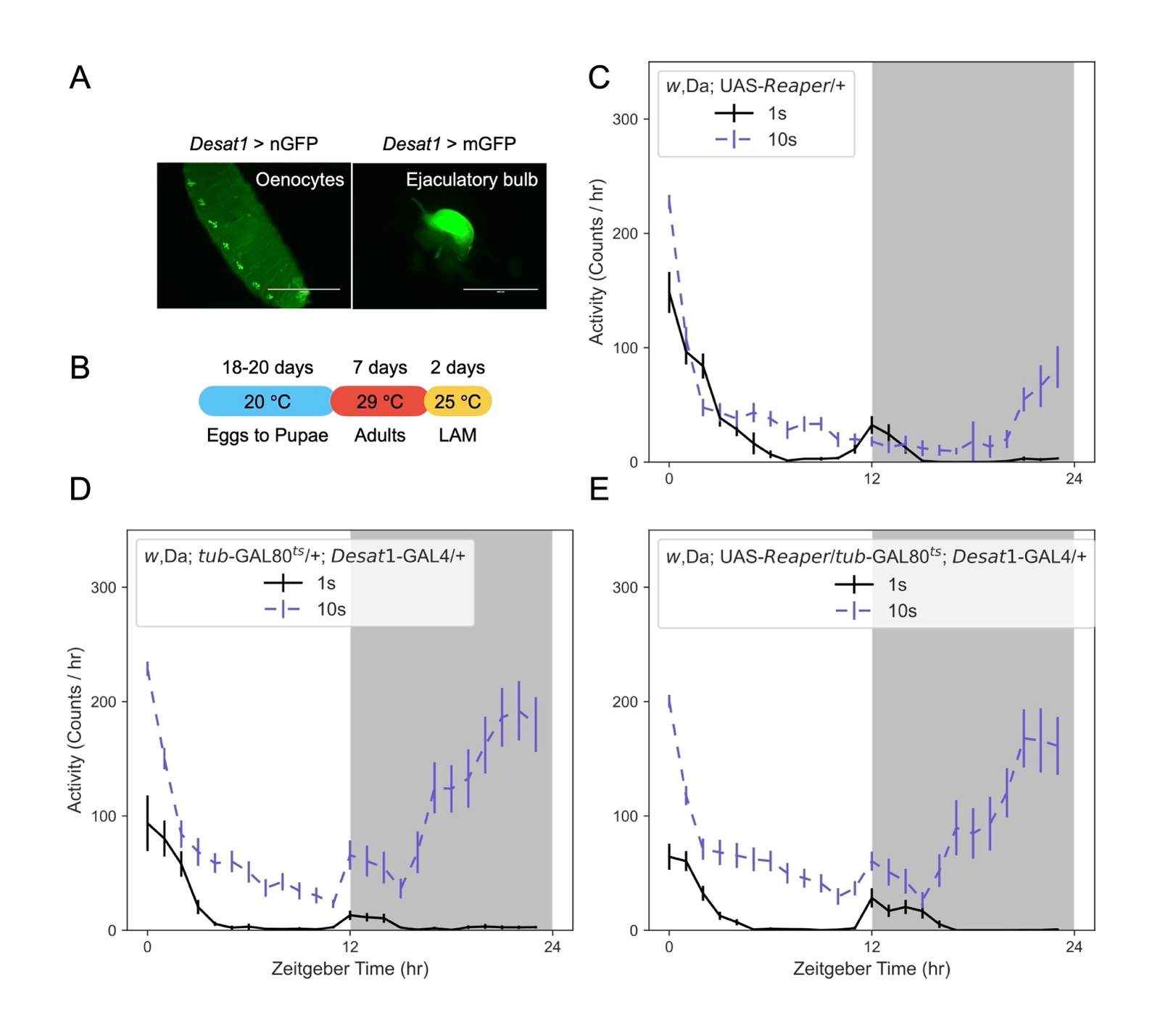

**Figure S7: Disrupting adult cuticular hydrocarbon synthesis does not affect group hyperactivity.**

(A) *Desat1-*GAL4 drives nuclear nGFP expression in oenocytes, and membrane-bound mGFP in the ejaculatory bulb. Scale bars: 1000 μm (left) or 400 μm (right). (B) Scheme for thermogenetic ablation of oenocytes followed by behavioral testing. (C-E) Activity over 24 hr of males with adult-specific ablation of oenocytes (E) compared to controls harboring either the *Reaper* (C) or *GAL4* (D) transgenes. Disruption of cuticular hydrocarbon synthesis did not disrupt group hyperactivity. N = 16-24 vials per condition.
