## Supplemental Table 1 for "Chemosensation drives divergent social behavior in *Drosophila*"

**Supplemental Table 1. Neuronal screen for altered nighttime group hyperactivity**

| **X (gene)** | **Nighttime group hyperactivity (10s/1s)^a^** | | |
| --- | --- | --- | --- |
|  | UAS*-Reaper* / *+* | *Or-X-*GAL4 / *+* | *Or-X-*GAL4 > *Reaper* |
| *Orco* | 2.6 | 4.8 | 1.2^b^ |
| *Or10a* | 4.9 | 11.1 | 4.5 |
| *Or13a* | 3.1 | 4.2 | 4.8 |
| *Or19a* | 1.0 | 3.5 | 5.8 |
| *Or22a* | 2.1 | 0.9 | 1.7 |
| *Or33a* | 1.0 | 1.8 | 1.2 |
| *Or33b* | 5.3 | 11.5 | 5.2 |
| *Or35a* | 2.4 | 5.3 | 2.3 |
| *Or43a* | 4.4 | 2.7 | 1.3 |
| *Or43a* (2nd trial) | 17.9 | 11.4 | 3.5 |
| *Or47a* | 4.4 | 33.8 | 2.8 |
| *Or56a* | 2.4 | 3.4 | 2.0 |
| *Or59b* | 3.1 | 2.1 | 2.0 |
| *Or65c* | 4.9 | 3.1 | 3.2 |
| *Or67b* | 4.4 | 5.7 | 2.2 |
| *Or7a* | 2.1 | 1.7 | 2.3 |
| *Or85a* | 1.3 | 12.1 | 1.5 |
| *Or85a* (2nd trial) | 4.4 | 8.3 | 3.4 |
| *Or85a* (3rd trial) | 1.5 | 11.1 | 2.0 |
| *Or85b* | 1.5 | 2.5 | 2.0 |
| *Or85f* | 2.1 | 2.0 | 1.8 |
| *Or9a* | 4.3 | 2.5 | 2.9 |
| *Or67a* | 4.0 | 9.6 | 2.8 |
| *Or43b* | 6.7 | 8.4 | 4.0 |
|  | UAS*-Kir2.1* / *+* | *Or-X-*GAL4 / *+* | *Or-X-*GAL4 > *Kir2.1* |
| *Orco* | 5.2 | 3.2 | 0.6 |
| *Or43a* | 5.2 | 3.6 | 2.5 |
| *Or67a* | 1.5 | 1.7 | 5.2 |
| *Orco* (2nd trial) | 2.5 | 2.4 | 1.1 |
| *Or43a* (2nd trial) | 2.5 | 6.3 | 1.3 |
| *Or67b* | 3.8 | 2.8 | 1.4 |
| *Or47a* | 3.8 | 3.4 | 2.3 |
| *Orco* (3rd trial) | 2.6 | 2.2 | 0.7 |
| *Or43a* (3rd trial) | 1.5 | 2.2 | 0.9 |
| *Or19a* | 3.3 | 2.6 | 2.8 |
| *Or67a* (2nd trial) | 1.3 | 1.9 | 1.5 |
|  | *Or-X-*GAL4 > *impTNT* | *Or-X-*GAL4 > *TNT* |  |
| *Orco* | 2.5 | 0.5^c^ |  |
| *Or43a* | 5.8 | 1.3 |  |
| *Orco* (2nd trial) | 2.3 | 1.1 |  |
| *Or43a* (2nd trial) | 4.1 | 1.1 |  |
| *Orco* (3rd trial) | 1.7 | 0.6 |  |
| *Or43a* (3rd trial) | 11.8 | 1.9 |  |

^a^Response to group-housing expressed as a ratio of activity per fly in groups vs. solo-housed from the 12-hr dark period. Males used in all experiments (N = 6-8 vials per condition).

^b^Underlined values are significantly different from both transgenic controls (*p* < 0.05, Dunn’s test with Bonferroni correction).

^c^Underlined values are significantly different than the inactive *TNT* (*impTNT*) control (Mann-Whitney U test).
