## Supplemental Table 2 for "Chemosensation drives divergent social behavior in *Drosophila*"

**Supplemental Table 2. Effect of genetic variation on nighttime locomotor activity**

| **DGRP lines^a^** | | **Beam break counts^b^** | | **Ratio^c^** |
| --- | --- | --- | --- | --- |
| BDSC # | Line # | Solo (1s) | Group (10s) | 10s/1s |
| 28122 | 21 | 783.3 | 1565.2 | 2.0 |
| 28123 | 26 | 68.3 | 224.6 | 3.3 |
| 29651 | 40 | 1215.0 | 1892.7 | 1.6 |
| 28128 | 45 | 364.7 | 481.1 | 1.3 |
| 29652 | 57 | 221.0 | 234.2 | 1.1 |
| 28129 | 59 | 155.5 | 365.2 | 2.3 |
| 28132 | 75 | 528.2 | 385.6 | 0.7 |
| 28134 | 83 | 642.5 | 330.6 | 0.5 |
| 28274 | 85 | 154.3 | 263.5 | 1.7 |
| 28136 | 91 | 2113.8 | 2276.5 | 1.1 |
| 28137 | 93 | 442.5 | 385.4 | 0.9 |
| 28138 | 101 | 1467.3 | 536.8 | 0.4 |
| 28140 | 109 | 138.5 | 175.9 | 1.3 |
| 28142 | 136 | 9.0 | 123.2 | 13.7 |
| 28144 | 142 | 429.7 | 525.0 | 1.2 |
| 28145 | 149 | 311.5 | 1079.0 | 3.5 |
| 28146 | 153 | 1391.3 | 591.6 | 0.4 |
| 28147 | 158 | 106.0 | 148.5 | 1.4 |
| 28150 | 177 | 913.0 | 1172.6 | 1.3 |
| 28151 | 181 | 37.7 | 759.3 | 20.2 |
| 28152 | 189 | 132.8 | 341.2 | 2.6 |
| 28153 | 195 | 113.3 | 123.1 | 1.1 |
| 25174 | 208 | 720.8 | 1593.5 | 2.2 |
| 28154 | 217 | 191.0 | 1390.9 | 7.3 |
| 28157 | 228 | 352.0 | 856.3 | 2.4 |
| 29653 | 229 | 382.7 | 81.3 | 0.2 |
| 28275 | 235 | 43.7 | 354.2 | 8.1 |
| 28160 | 237 | 422.2 | 979.1 | 2.3 |
| 28161 | 239 | 457.3 | 527.9 | 1.2 |
| 28164 | 280 | 689.5 | 702.0 | 1.0 |
| 28165 | 287 | 493.8 | 779.1 | 1.6 |
| 25175 | 301 | 1206.2 | 640.9 | 0.5 |
| 25176 | 303 | 299.5 | 60.7 | 0.2 |
| 25177 | 304 | 82.8 | 144.4 | 1.7 |
| 37525 | 306 | 332.2 | 41.0 | 0.1 |
| 25179 | 307 | 179.0 | 56.0 | 0.3 |
| 28166 | 309 | 132.7 | 335.3 | 2.5 |
| 28167 | 317 | 1182.3 | 1001.7 | 0.8 |
| 28168 | 318 | 80.2 | 256.4 | 3.2 |
| 29654 | 320 | 183.3 | 257.0 | 1.4 |
| 29655 | 321 | 914.7 | 607.1 | 0.7 |
| 25182 | 324 | 197.2 | 162.5 | 0.8 |
| 25183 | 335 | 2.3 | 56.8 | 24.3 |
| 28172 | 336 | 43.5 | 207.2 | 4.8 |
| 28173 | 338 | 232.8 | 326.5 | 1.4 |
| 28174 | 340 | 773.2 | 489.8 | 0.6 |
| 28176 | 350 | 399.8 | 392.2 | 1.0 |
| 28177 | 352 | 1440.7 | 2158.2 | 1.5 |
| 28178 | 356 | 2147.8 | 1238.5 | 0.6 |
| 25184 | 357 | 963.7 | 446.5 | 0.5 |
| 25185 | 358 | 768.8 | 840.9 | 1.1 |
| 28179 | 359 | 199.4 | 258.8 | 1.3 |
| 25186 | 360 | 336.0 | 389.5 | 1.2 |
| 28180 | 361 | 407.5 | 2499.1 | 6.1 |
| 25187 | 362 | 518.3 | 430.4 | 0.8 |
| 25445 | 365 | 596.2 | 818.8 | 1.4 |
| 28182 | 370 | 532.5 | 842.5 | 1.6 |
| 28183 | 371 | 577.2 | 1417.2 | 2.5 |
| 28184 | 373 | 103.3 | 339.2 | 3.3 |
| 28185 | 374 | 1188.3 | 1248.0 | 1.1 |
| 25188 | 375 | 654.0 | 774.4 | 1.2 |
| 28186 | 377 | 1339.7 | 1364.7 | 1.0 |
| 25189 | 379 | 221.7 | 90.2 | 0.4 |
| 25190 | 380 | 194.7 | 1276.5 | 6.6 |
| 28189 | 382 | 235.7 | 225.2 | 1.0 |
| 28191 | 385 | 433.7 | 296.3 | 0.7 |
| 28192 | 386 | 1852.5 | 1431.7 | 0.8 |
| 25191 | 391 | 839.3 | 2138.2 | 2.5 |
| 28194 | 392 | 1147.3 | 1522.5 | 1.3 |
| 25192 | 399 | 1323.5 | 595.4 | 0.4 |
| 29656 | 405 | 237.5 | 1131.3 | 4.8 |
| 28278 | 409 | 159.8 | 354.1 | 2.2 |
| 28196 | 426 | 394.3 | 883.1 | 2.2 |
| 25193 | 427 | 629.8 | 232.6 | 0.4 |
| 25194 | 437 | 191.8 | 1636.2 | 8.5 |
| 29658 | 439 | 74.2 | 241.6 | 3.3 |
| 28197 | 440 | 246.3 | 1492.2 | 6.1 |
| 28198 | 441 | 85.2 | 475.9 | 5.6 |
| 28199 | 443 | 1217.8 | 198.6 | 0.2 |
| 28202 | 491 | 192.3 | 468.8 | 2.4 |
| 28203 | 492 | 527.2 | 159.0 | 0.3 |
| 28204 | 502 | 279.8 | 1286.3 | 4.6 |
| 28205 | 508 | 1136.3 | 551.5 | 0.5 |
| 25197 | 517 | 867.2 | 1522.3 | 1.8 |
| 29660 | 530 | 575.0 | 623.1 | 1.1 |
| 28208 | 535 | 415.0 | 2006.9 | 4.8 |
| 25198 | 555 | 318.2 | 260.9 | 0.8 |
| 28211 | 563 | 2567.3 | 777.7 | 0.3 |
| 28212 | 584 | 546.7 | 178.9 | 0.3 |
| 28213 | 589 | 1532.8 | 401.4 | 0.3 |
| 28215 | 595 | 870.3 | 1187.8 | 1.4 |
| 28218 | 703 | 2845.0 | 584.4 | 0.2 |
| 25200 | 707 | 98.2 | 335.8 | 3.4 |
| 25201 | 712 | 1038.5 | 2080.5 | 2.0 |
| 28219 | 716 | 277.0 | 507.0 | 1.8 |
| 28220 | 721 | 121.8 | 680.7 | 5.6 |
| 25203 | 732 | 1115.7 | 1253.0 | 1.1 |
| 28222 | 737 | 561.7 | 1217.6 | 2.2 |
| 28223 | 738 | 589.8 | 852.7 | 1.4 |
| 28227 | 761 | 1606.5 | 145.1 | 0.1 |
| 25204 | 765 | 1306.3 | 583.5 | 0.4 |
| 28229 | 776 | 331.3 | 2124.9 | 6.4 |
| 28230 | 783 | 545.7 | 810.4 | 1.5 |
| 25206 | 786 | 18.8 | 105.0 | 5.6 |
| 28231 | 787 | 345.3 | 263.4 | 0.8 |
| 28233 | 796 | 2054.2 | 454.4 | 0.2 |
| 25207 | 799 | 671.7 | 579.8 | 0.9 |
| 28234 | 801 | 281.2 | 798.0 | 2.8 |
| 28235 | 802 | 527.5 | 352.0 | 0.7 |
| 28236 | 804 | 1254.3 | 1296.3 | 1.0 |
| 28238 | 808 | 1402.0 | 470.0 | 0.3 |
| 28239 | 810 | 337.0 | 366.5 | 1.1 |
| 28241 | 818 | 550.0 | 625.2 | 1.1 |
| 25208 | 820 | 3071.0 | 1230.6 | 0.4 |
| 28243 | 821 | 149.3 | 446.0 | 3.0 |
| 28244 | 822 | 2359.8 | 279.1 | 0.1 |
| 28245 | 832 | 1027.8 | 633.6 | 0.6 |
| 28246 | 837 | 22.3 | 491.1 | 22.0 |
| 28247 | 843 | 69.2 | 58.2 | 0.8 |
| 28249 | 850 | 325.2 | 494.7 | 1.5 |
| 28250 | 853 | 550.7 | 1093.3 | 2.0 |
| 28251 | 855 | 530.5 | 656.5 | 1.2 |
| 28252 | 857 | 728.7 | 2004.2 | 2.8 |
| 25210 | 859 | 474.7 | 450.8 | 0.9 |
| 28253 | 861 | 71.3 | 724.6 | 10.2 |
| 28254 | 879 | 138.8 | 683.1 | 4.9 |
| 28255 | 882 | 1018.2 | 830.8 | 0.8 |
| 28257 | 890 | 815.0 | 1339.3 | 1.6 |
| 28258 | 892 | 968.7 | 876.0 | 0.9 |
| 28260 | 897 | 493.7 | 255.8 | 0.5 |
| 28261 | 900 | 388.5 | 148.7 | 0.4 |
| 28262 | 907 | 235.7 | 307.7 | 1.3 |
| 28263 | 908 | 59.3 | 133.4 | 2.2 |
| 28264 | 911 | 786.5 | 532.8 | 0.7 |
| 28265 | 913 | 739.2 | 850.2 | 1.2 |

^a^DGRP line number and the corresponding Bloomington (BDSC) stock ID number.

^b^Underlined Average nighttime beam breaks per fly from group- or solo-housing. Males used in all experiments (N = 6 vials per condition).

^c^Response to group-housing expressed as a ratio of activity per fly in groups vs. solo-housed.
